# Biallelic variants in LARS1 induce steatosis in developing zebrafish liver via enhanced autophagy

**DOI:** 10.1101/2023.09.21.558924

**Authors:** Masanori Inoue, Wulan Apridita Sebastian, Shota Sonoda, Hiroaki Miyahara, Nobuyuki Shimizu, Hiroshi Shiraishi, Miwako Maeda, Kumiko Yanagi, Tadashi Kaname, Reiko Hanada, Toshikatsu Hanada, Kenji Ihara

**Affiliations:** Department of Pediatrics, Oita University Faculty of Medicine, Oita, Japan; Department of Neuropathology, Institute for Medical Science of Aging, Aichi Medical University, Aichi, Japan; Department of Cell Biology, Oita University Faculty of Medicine, Oita, Japan; Department of Genome Medicine, National Center for Child Health and Development, Tokyo, Japan; Department of Neurophysiology, Oita University Faculty of Medicine, Oita, Japan

## Abstract

Acute liver failure is a life-threatening condition during infancy. Biallelic pathogenic variants in *LARS1* cause infantile liver failure syndrome type 1 (ILFS1), which is characterized by acute hepatic failure in infants. LARS functions as a protein associated with mTORC1 and plays a crucial role in amino acid-triggered mTORC1 activation and autophagy regulation. A previous study demonstrated that *larsb*-knockout zebrafish show a condition resembling ILFS. However, a comprehensive analysis of *larsb*-knockout zebrafish has not yet been performed because of early mortality. We herein generated a long-term viable zebrafish model carrying a *LARS1* variant identified in an ILFS1 patient (*larsb-I451*F zebrafish) and analyzed the pathogenesis of the affected liver of ILFS1. Hepatic dysfunction is most prominent in ILFS1 patients during infancy; correspondingly, the *larsb-I451F* zebrafish manifested hepatic anomalies during the developmental stages. The *larsb-I451F* zebrafish demonstrates augmented lipid accumulation within the liver under autophagy activation. Inhibition of DGAT1, which converts fatty acids to triacylglycerols, improved lipid droplets in the liver of *larsb-I451F* zebrafish. Notably, treatment with an autophagy inhibitor ameliorated hepatic lipid accumulation in this model. Our findings suggested that enhanced autophagy caused by biallelic *LARS1* variants contributes to ILFS1-associated hepatic dysfunction. Furthermore, the *larsb-I451F* zebrafish model, which has a prolonged survival rate compared to the *larsb*-knockout model, highlights its potential utility as a tool for investigating the pathophysiology of ILFS1-associated liver dysfunction.

**Author Summary:** Infantile liver failure (ALF) is a rare but life-threatening condition primarily caused by various genetic and infectious factors during infancy. Comprehensive research into its causes is crucial for treatment decisions, including liver transplantation and supportive interventions. While specific therapies exist for some conditions, a significant proportion of infant ALF cases remains unresolved. Recent advances in genetic sequencing have identified congenital disorders, particularly involving the *LARS1* gene, as contributors to ALF. *LARS1* is essential for regulating processes related to amino acids and autophagy. To better understand this condition, we created a zebrafish model carrying specific *LARS1* gene variants seen in ALF patients. These zebrafish displayed liver abnormalities similar to those observed in infants with ALF. Our study revealed that enhanced autophagy, triggered by biallelic *LARS1* variants, plays a significant role in liver dysfunction associated with ALF. Notably, inhibiting specific enzymes involved in fat metabolism and autophagy showed promising results in reducing hepatic lipid accumulation in our zebrafish model. This research provides insights that may lead to improved understanding and potential treatments for this devastating condition.

## Introduction

Acute liver failure (ALF) in infancy is a rare but life-threatening event [1]. The primary disorders causing ALF during this period include hereditary metabolic disorders, such as mitochondrial respiratory chain disorders, type I hereditary tyrosinemia, and urea cycle disorders [2]. Congenital infections of viruses or bacteria, such as cytomegalovirus, toxoplasma, or herpes, and gastrointestinal alloimmune diseases, such as neonatal hemochromatosis, are also known to cause ALF [2, 3]. Other types of ALF result from hyperimmune activation under a genetic predisposition to cholestasis, such as hemophagocytic syndrome or Niemann-Pick disease type C [3, 4].

Given the above, conducting a comprehensive investigation into the etiology of ALF in infancy is crucial for treatment decisions, including liver transplantation, along with supportive care with dietary therapy or supplementary intervention [5–8]. Disease-specific treatments have been established for some diseases, such as chemotherapy for hemophagocytic syndrome, inhibitors of glycosphingolipid synthesis, miglustat for Niemann-Pick disease type C, inhibitor of tyrosine degradation, and nitisinone (NTBC) [5, 9, 10]. Nevertheless, a significant proportion of infant ALF cases (approximately 20%-50%) remain unresolved [1, 2, 11].

Recent advances in whole-exome sequencing (WES) have revealed new congenital disorders that cause ALF in infants. Since 2012, congenital defects in aminoacyl-tRNA synthetases (ARSs) have been reported to cause ALF [12–17]. ARSs are essential enzymes that catalyze the ligation of amino acids with their cognate transfer RNAs, which is the first step in protein synthesis [18–21]; For example, leucyl-tRNA synthetase (LARS) catalyzes the ligation of leucine to leucine tRNA. Biallelic pathogenic variants in the *LARS1* gene lead to infantile hepatopathy, recently known as infantile liver failure syndrome type 1 (ILFS1), which is characterized by ALF within the first few months after birth. It is also associated with failure to thrive, anemia, microcephaly, muscular hypotonia, and seizures [14, 22]. LARS plays a unique, non-canonical role as a mammalian target of rapamycin complex 1 (mTORC1)-associated protein required for amino acid-induced mTORC1 activation, which acts as an intracellular leucine sensor for mTORC1 signaling [23–26]. Thus, LARSs play broad roles in cellular homeostasis, including translation control, transcriptional regulation, tumorigenesis, and senescence [23–28].

Our previous research using *larsb*-knockout zebrafish demonstrated that mutant zebrafish exhibited a phenotype similar to that of ILFS1 [29]. Excessive autophagy activation was observed in *larsb*-knockout zebrafish, and the suppression of autophagy by bafilomycin treatment significantly recovered the liver size and improved the survival curve [29]. However, early lethality, probably due to severe liver damage, nervous system disorders, and anemia in *larsb*-knockout larvae, did not allow us to analyze the exact molecular mechanism by which LARS pathogenic variants affect the development and function of the liver in ILFS1 patients.

To further evaluate the role of LARS and the effects of its defect in the pathogenesis of the liver, we generated *larsb*-knockin zebrafish with a biallelic missense variant of the *LARS1* gene identified in the ILFS1 patient in our hospital. We then investigated the molecular function of Lars in the context of ILFS1 pathogenesis.

## Results

### ILFS1 patient with liver dysfunction

The patient was the first male child born to a non-consanguineous Japanese couple, and his younger brother and parents had no congenital abnormalities, including liver disease (Fig 1A). He was delivered at 37 weeks’ gestation with a birth weight of 2,320 g (9.2 %tile). Marked hepatomegaly and failure to thrive were detected during routine checkups by a primary pediatrician at seven months old, and he was referred to our hospital. At 8 months old, his height was 62.2 cm (-3.3 standard deviations [SD]), his body weight was 6.6 kg (-2.1 SD), and his head circumference was 43.9 cm (0.0 SD). He presented with a cherubic face with full cheeks, hepatomegaly (approximately 8 cm below the costa), and mild hypotonia. He was able to control his head by himself but lacked the ability to roll over and sit up unaided. Abdominal computed tomography revealed a diffuse, low-density, enlarged liver (Fig 1B).

Laboratory findings demonstrated mild elevation of serum AST and ALT levels (103 U/l and 70 U/l, respectively) with mild anemia (hemoglobin 10.6 g/dl). Several days later, he developed a high fever for the first time after birth, caused by a human herpesvirus 6 infection. His liver dysfunction soon progressed to ALF as elevation of transaminases (AST 870 U/l, ALT 263 U/l) with reduction of protein synthesis (PT-INR 1.53) and hypoalbumininema (albumin 2.47 g/dl), remarkable anemia (hemoglobin 6.3 g/dl), and thrombocytopenia (platelet count 19,000/μl) (Fig 1C). He continued to have a fever, generalized edema, oliguria, and respiratory distress and received treatments that included acetaminophen administration, albumin infusion, red blood cell transfusion, and oxygen therapy.

His critical condition recovered with defervescence after five days. Following this episode, he experienced four episodes of febrile illnesses, including acute pharyngitis, hand-foot-mouth disease, and acute gastroenteritis, over the next two years. However, the symptoms appeared to be mild, and ALF did not recur, as transaminase levels peaked at AST 80-220 U/l and ALT 70-260 U/l during these episodes, and growth retardation gradually normalized by 3 years old (Fig 1C and 1D). His febrile episodes after his first three years of life included negligible deterioration of the liver function. His psychomotor development progressed normally, with a developmental quotient at 3 years old as assessed by the Enjohji Developmental Test in Infancy and Early Childhood of 106; however, cognitive dysfunction was identified at 6 years old using the Wechsler Intelligence Scale for Children-Fourth edition.

The patient is now 12 years old, and the most recent data are as follows: height, 147.3 cm (-0.2 SD); weight, 34.9 kg (body mass index 16.1); serum AST level, 24 U/l; serum ALT level, 22 IU/l; serum albumin level 4.04 g/dl; hemoglobin 12.6 g/dl; platelet count, 395,000/μl; and white blood cell count, 6,870/μl, indicating a normal physical growth and liver function with mild anemia.

**Fig 1.**
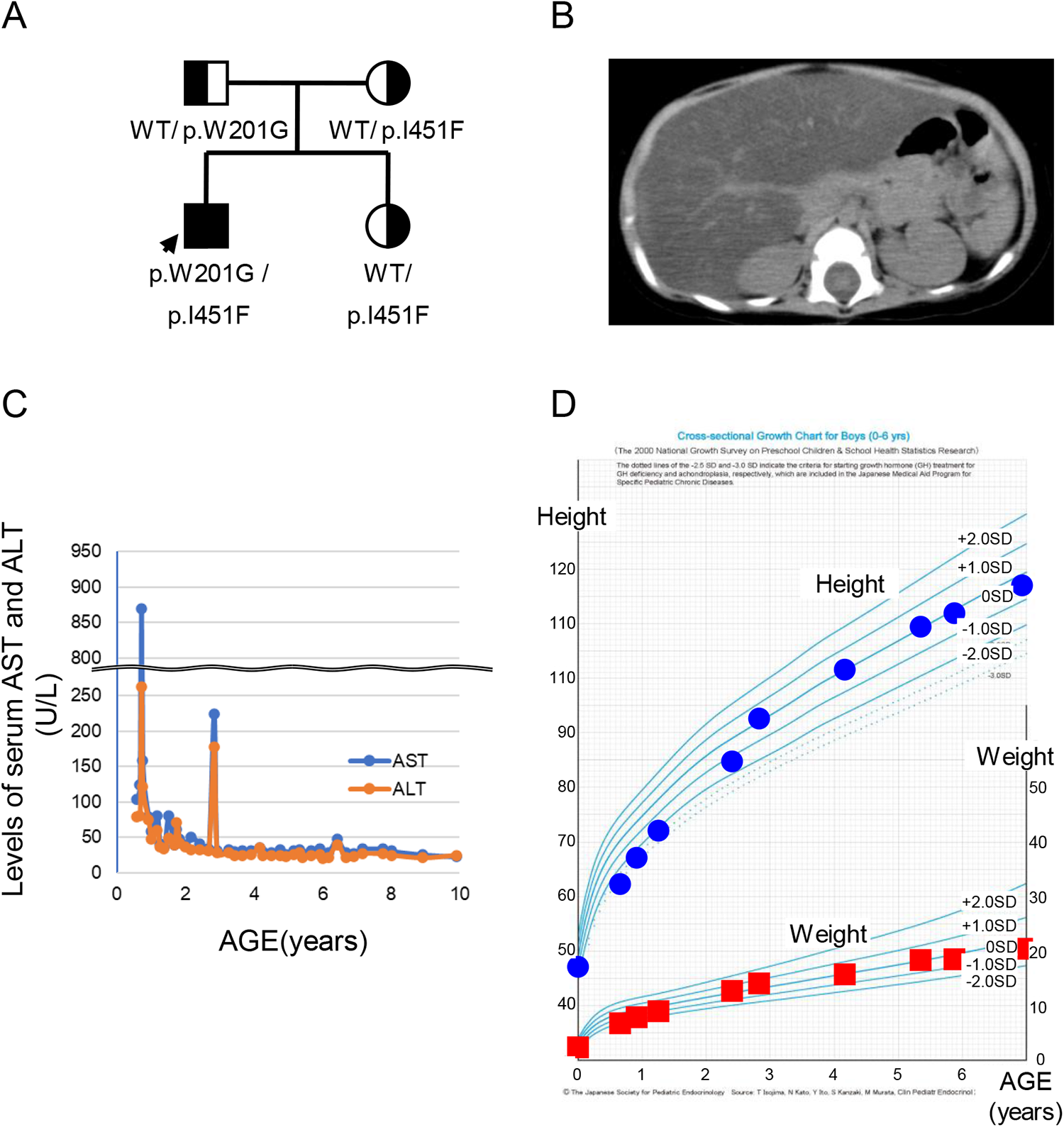
Clinical information of an infantile liver failure syndrome type 1 patient with biallelic LARS1 variants in our hospital. (A) Pedigree of the family. (B) Abdominal computed tomography image at eight months old. (C) Changes in serum levels of AST and ALT. (D) Developmental curve.

### *LARS1* as a single candidate gene by WES

WES using the child-parent trio revealed compound heterozygosity in the infant for two potentially pathogenic variants of *LARS1* [NM_020117.9] (Fig 1A). One missense variant, c.601T>G; p.W201G in exon 7 [NM_020117.9], was paternally inherited and had not been previously reported in ClinVar (https://www.ncbi.nlm.nih.gov/clinvar). An *in silico* analysis suggested that W201G probably damaged the protein structure and/or function (Polyphen2: score 1.000; probably damaging [http://genetics.bwh.harvard.edu/pph2/]) (S1 Table). Another missense variant, c.1351A>T; p.I451F in exon 14 [NM_020117.9], was maternally inherited and had been previously described in a Japanese patient with ILFS1 [30]. It is located in the LARS editing domain (Fig 2A). Importantly, 9 of the 23 pathogenic variants previously reported in ILFS1 patients were located in this domain. Three editing-domain variants, including I451F, showed severe symptoms during the neonatal period [14, 22, 30–33]. The *in silico* analysis predicted that I451F was also probably damaging to the protein structure and/or function (PolyPhen-2:0.921; probably damaging) (S1 Table). Notably, both missense variants affected the evolutionarily conserved residues (Fig 2B).

**Fig 2.**
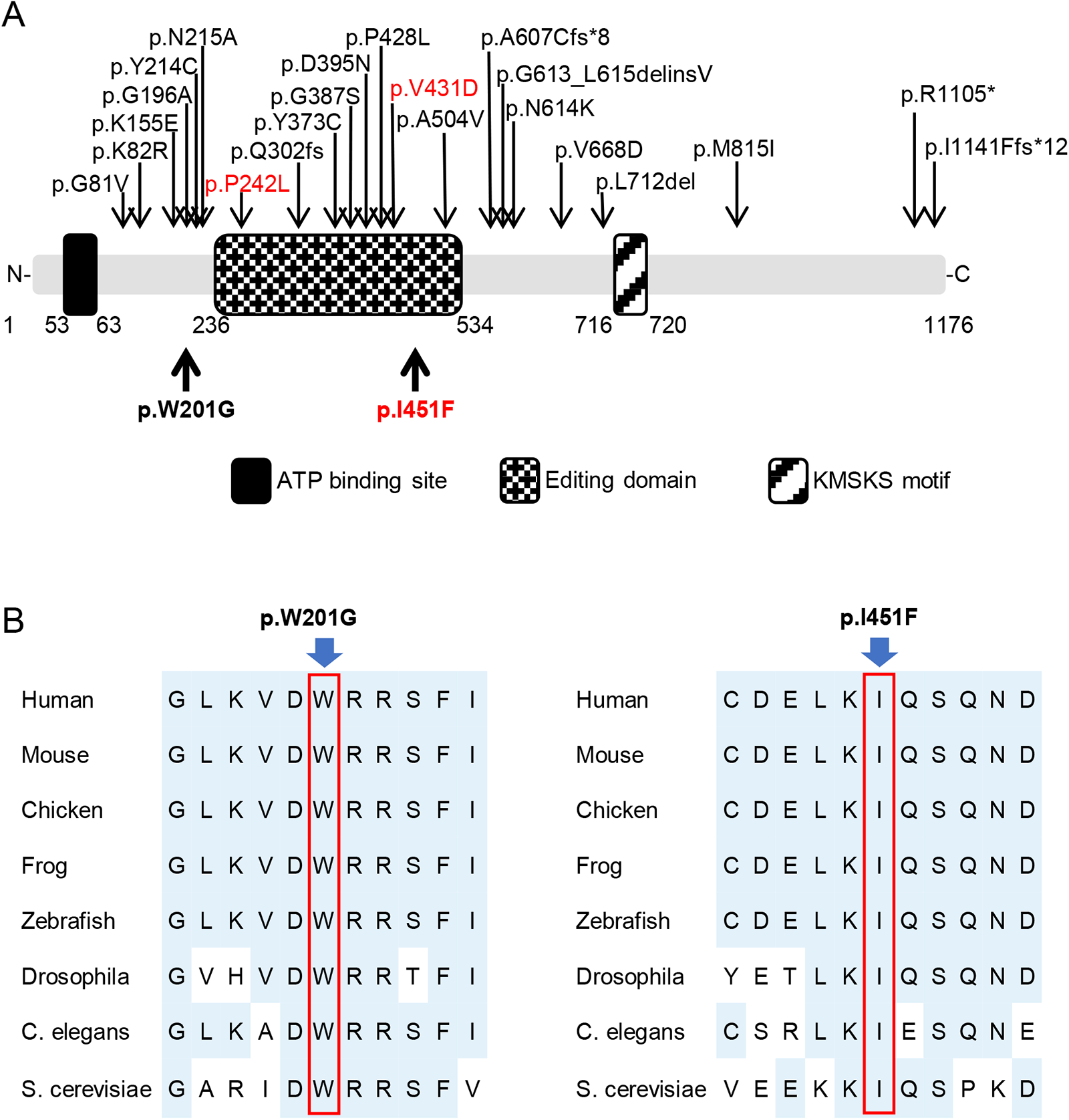
Leucine-tRNA synthetase (LARS) mutations. (A) LARS domains and pathogenic variants found in infantile liver failure syndrome type 1 patients. Variants in our patients are shown in bold. Variants in another reported case with severe manifestation in the neonatal period are in red. (B) Conservations of the missense variant in LARS.

### Liver defects in *larsb-I451F* zebrafish during liver development

To assess the pathological relevance of LARS variants in the liver, we generated A-to-T at codon 1351 and C-to-T at codon 1353 knock-in zebrafish lines using CRISPR/Cas9. To obtain more efficient knock-in using genome editing, we replaced the two bases that changed the PAM sequence (Fig 3A). Among the pathogenic variants in the *LARS* gene (p.W201G/p.I451F) identified in our patient, we focused on the p.I451F variant, as it has been found in other Japanese patients, suggesting a Japanese founder effect, and is located within the editing domain of the LARS protein, where pathological variants have accumulated [30]. We designed a model of the *larsb I451F* mutation (*larsb-I451F*) to elucidate the pathogenesis of ILFS1 (Fig 3B).

First, we measured the amount of Lars protein in the whole body of *larsb I451F* zebrafish larvae. Western blotting confirmed that the amount of Lars protein in *larsb-I451F* zebrafish was similar to that in wild-type (WT) *larsb* zebrafish (S1A and 1B Fig). Patients with ILFS1 exhibit hepatomegaly and liver damage with rapid progression after viral infection during neonatal and infancy [22]. To analyze the morphology of the liver, *larsb-I451F* zebrafish were crossed with Tg[*fabp10*:mcherry] transgenic zebrafish, which constitutively express mCherry fluorescent protein in the liver [34, 35]. Because zebrafish livers mature at the larval stage by five days old [36], we observed *larsb-I451F* zebrafish livers at approximately five days post-fertilization (dpf). At 5 dpf, *larsb-I451F* zebrafish exhibited increased liver circularity (Fig 3C and 3D), a common feature of liver diseases [37]. As *larsb-I451F* zebrafish grew, morphological abnormalities in the liver gradually improved by 7 dpf. In addition, *larsb-I451F* zebrafish survived to adulthood in the same manner as WT zebrafish. Since hepatoblasts proliferate and differentiate between 2 and 5 dpf in zebrafish livers [38], we found morphological abnormalities predominantly appearing in developing hepatocytes in *larsb-I451F* zebrafish.

**Fig 3.**
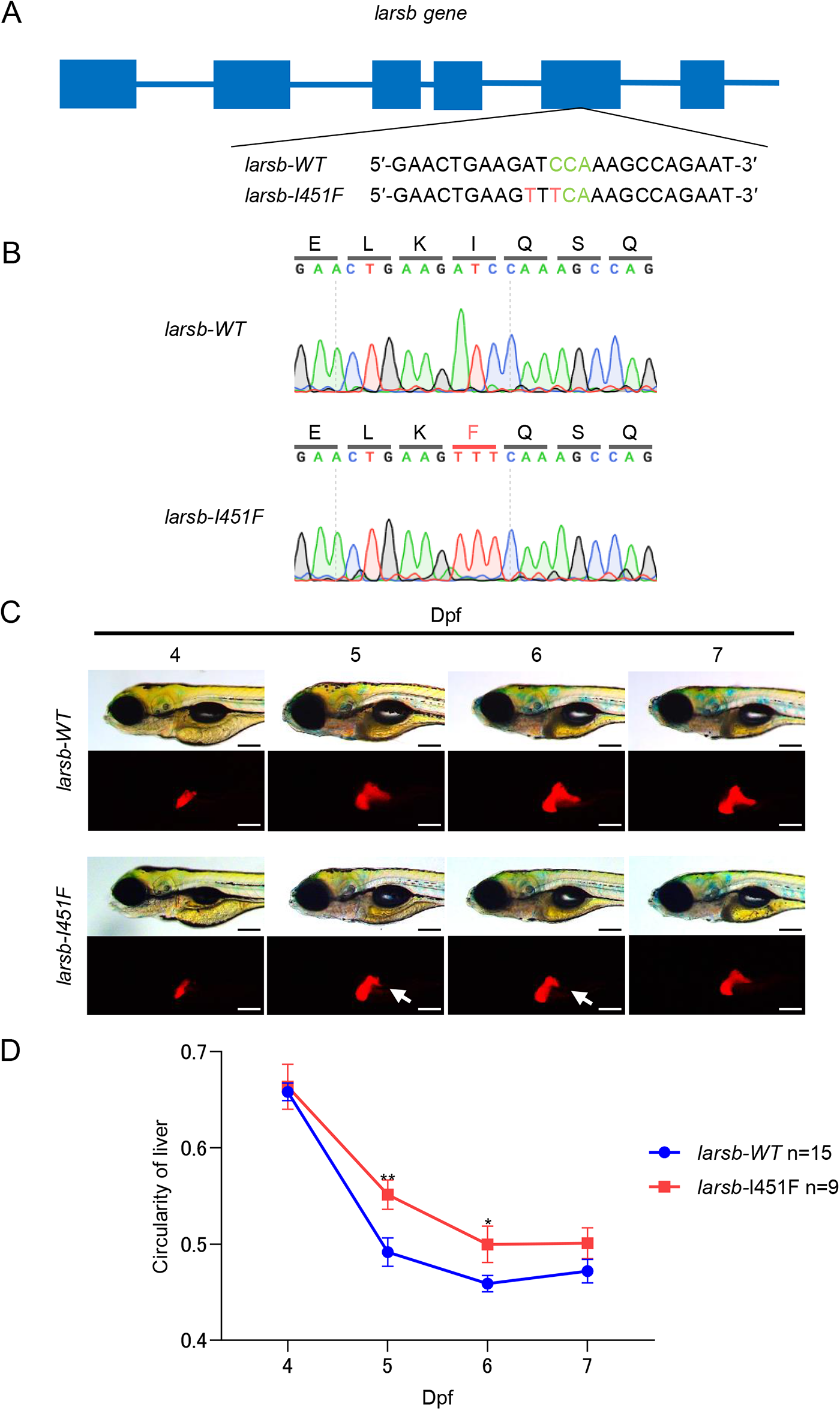
*Larsb*-knockin larvae display liver abnormality during the liver developmental stage. (A) Diagram showing the *larsb* genomic locus and *larsb*-knockin (*larsb-I451F*) zebrafish mutant genotype. (B) In the genomic sequencing analysis chromatograms, the mutation site in the *larsb-I451F* zebrafish is shown in red. (C) Morphological abnormality at 4 to 7 dpf in the livers of *larsb-I451F* larvae with a Tg[*fabp10*:mcherry] background. White arrows indicate the loss of liver edges in *larsb-I451F* larvae. Scale bar: 200 μm. (D) Circularity of liver in *larsb-I451F* larvae with a Tg[*fabp10*:mcherry] background (4 to 7 dpf). Error bars indicate SEM. *P < 0.05, **P < 0.01.

### Hepatic adiposity in *larsb-I451F* zebrafish

The liver was histopathologically analyzed. The livers of *larsb-I451F* larvae contained more vacuoles than those of *larsb-WT* larvae (Fig 4A and 4B). Multiple vacuoles in the cytoplasm and clear circular spaces with sharp outlines and contours are characteristic of fat-type vacuolation [39].

Next, to examine whether or not intrahepatic vacuoles in *larsb-I451F* zebrafish were lipid droplets, we evaluated intrahepatic lipids using fluorescent staining [40]. Many lipid droplets visualized by lipid dye droplet staining were observed in the livers of *larsb-I451F* larvae compared to those of *larsb-WT* larvae (Fig 4C and 4D). These data indicate that Lar dysfunction induces hepatic lipid droplet formation. Most patients with ILFS1 present with liver steatosis [22]. Thus, *larsb-I451F* zebrafish exhibited a phenotype analogous to that observed in ILFS1, indicating that the function of LARS in the liver is conserved between zebrafish and humans.

**Fig 4.**
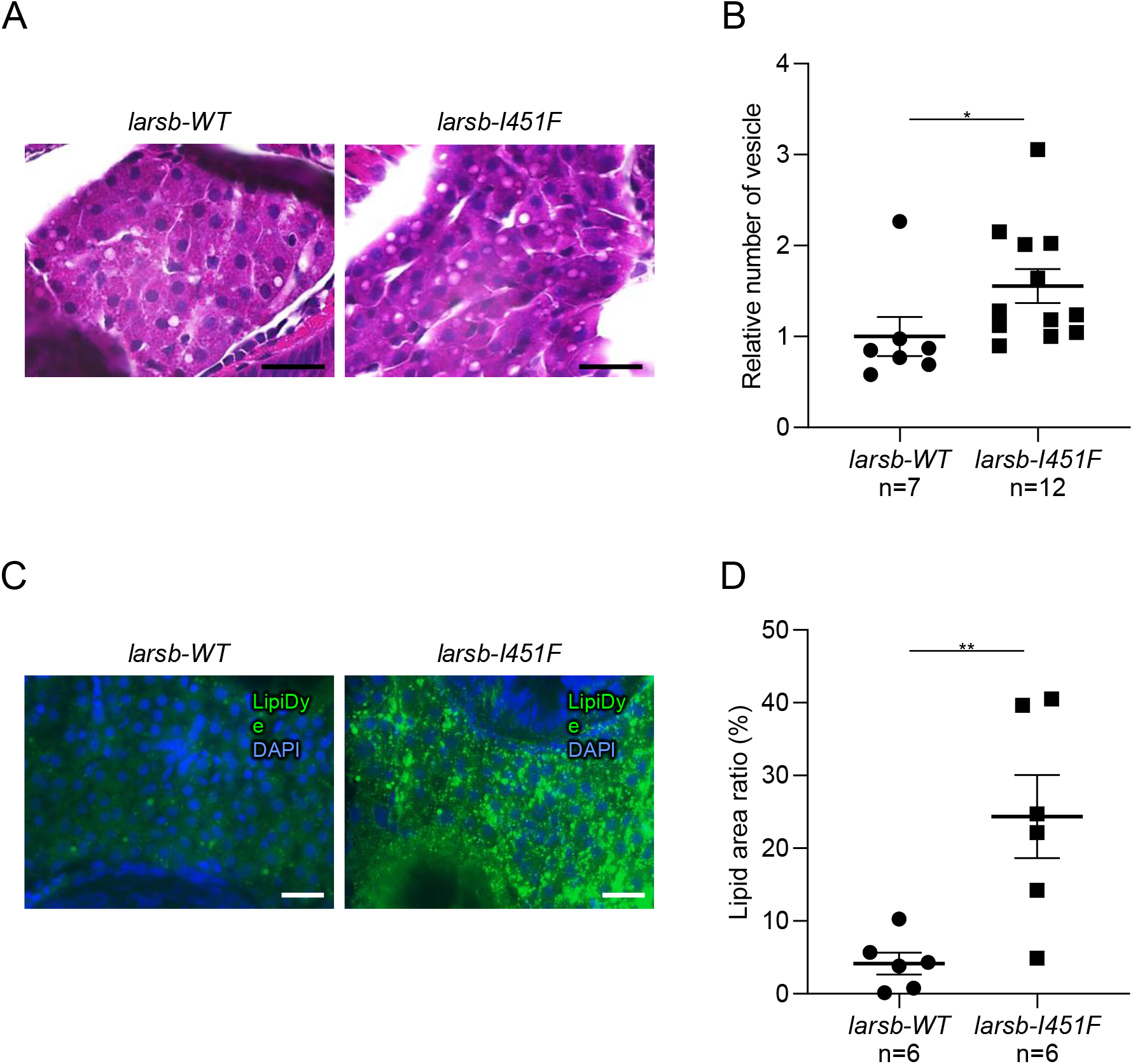
Histopathology and lipids staining of the liver in *larsb*-knockin larvae. (A) Hematoxylin and eosin staining of the liver in *larsb*-knockin (*larsb-I451F*) larvae at 5 dpf. Scale bar: 20 μm. (B) Quantification of the vesicle number in *larsb-I451F* larvae liver at 5 dpf. Error bars indicate SEM. *P < 0.05. (C) Lipids staining of the liver in *larsb-I451F* larvae at 5 dpf. Scale bar: 20 μm. (D) Quantification of the lipid area in *larsb-I451F* larvae liver at 5 dpf. Error bars indicate SEM. **P < 0.01. Dpf: days post fertilization.

### The *larsb-I451F* mutation augments autophagy in liver

Excessive activation of autophagy has been observed in *larsb*-deficient zebrafish [29]. Therefore, to assess whether or not autophagy is involved in liver abnormalities in *larsb-I451F* zebrafish, we evaluated the status of autophagy by fluorescent immunostaining for LC-3 and p62 in *larsb-I451F* larvae. LC-3, a downstream component of the autophagy pathway that participates in autophagosome formation, is widely used to monitor autophagy [41]. While the expression of p62, a selective autophagy substrate, did not differ markedly between *larsb-I451F* and *larsb-WT* larvae (S2A and 2B Fig), many autophagosomal structures visualized with LC-3 were observed in the livers of *larsb-I451F* larvae compared to *larsb-WT* larvae (Fig 5A and 5B). Therefore, Lar dysfunction appears to enhance autophagy in the developing liver.

Next, to validate whether or not the lipid droplets detected in the livers of *larsb-I451F* larvae were induced by enhanced autophagy, *larsb-I451F* larvae were treated with an inhibitor specific to diacylglycerol acyltransferase 1 (DGAT1) (A922500). DGAT1 and DGAT2 mediate the final step in the synthesis of triacylglycerols from fatty acids stored in lipid droplets [42, 43]. Because both DGAT1 and DGAT2 act on liver lipid droplet formation due to overnutrition, inhibition of DGAT1 alone does not usually improve lipid droplets [42–44]. In contrast, hepatic lipid accumulation via autophagy is specifically mediated by DGAT1 [44]. We demonstrated that A922500 treatment improved the accumulation of intrahepatic lipids in *larsb-I451F* larvae (S3A and 3B Fig), and consequently, it was likely that the accumulated lipid droplets in the livers of *larsb-I451F* zebrafish had been induced by autophagy.

To verify whether or not liver abnormalities in *larsb-I451F* larvae were due to excessive autophagy, we treated *larsb-I451F* larvae with the autophagy inhibitor bafilomycin A1. Bafilomycin treatment improved liver abnormalities and decreased liver circularity in *larsb-I451F* larvae at 5 dpf (Fig 5C and 5D). The accumulation of intrahepatic lipids was significantly reduced by bafilomycin treatment (Fig 5E and 5F). We concluded that hyperactivated autophagy induced by *larsb-I451F* was responsible for liver steatosis.

**Fig 5.**
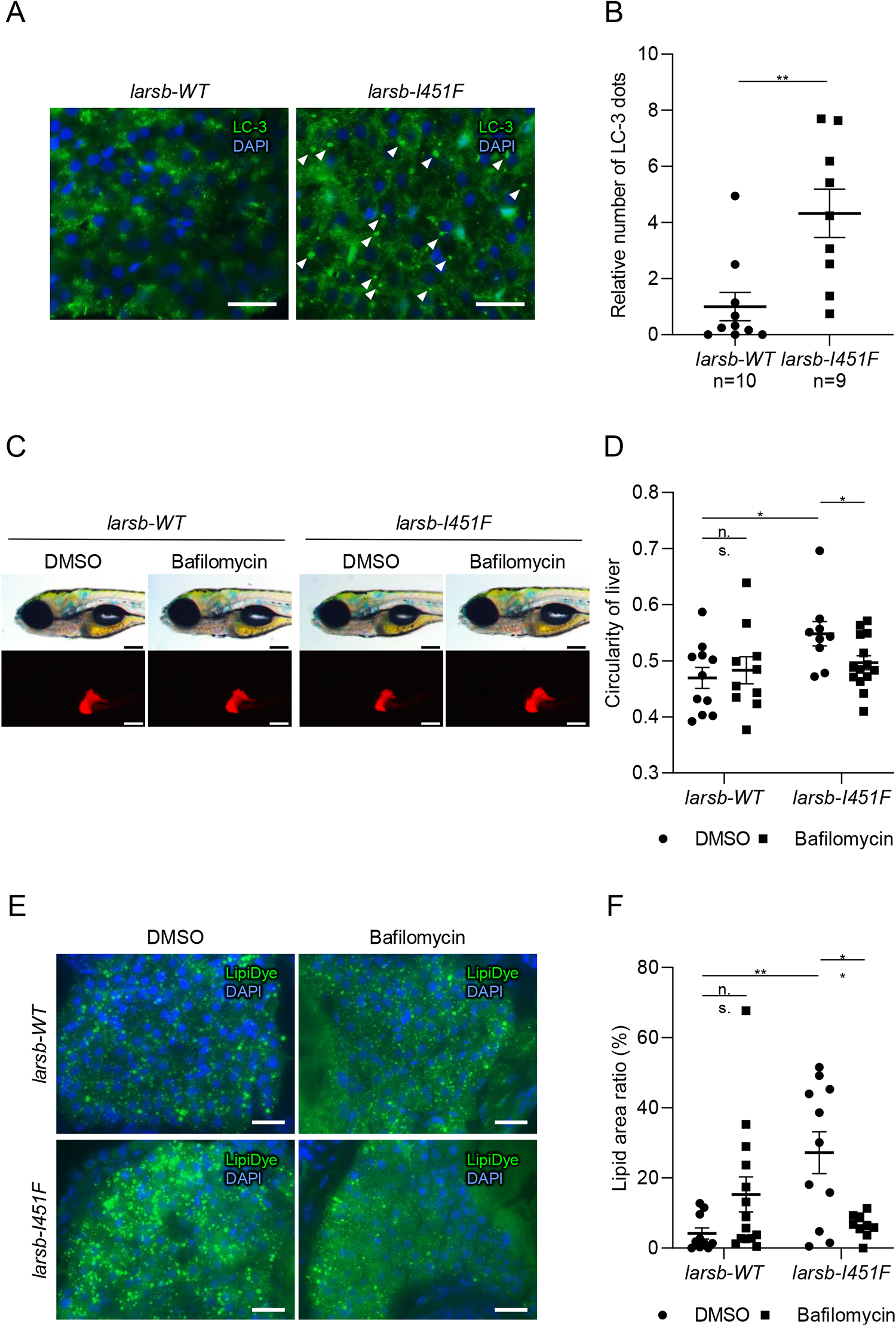
Enhanced autophagy in the liver of *larsb*-knockin larvae. (A) Immunostaining of LC-3 in the liver of *larsb*-knockin (*larsb-I451F*) larvae at 5 dpf. Scale bar: 20 μm. White arrowheads indicate LC3-positive dots. (B) Quantification of the number of LC-3 dots in *larsb-I451F* larvae liver at 5 dpf. Error bars indicate SEM. **P < 0.01. (C) Morphological abnormality in the livers of *larsb-I451F* larvae at 5 dpf with a Tg[*fabp10*:mcherry] background treated with DMSO or bafilomycin. Scale bar: 200 μm. (D) Circularity of the liver in *larsb-I451F* larvae at 5 dpf with a Tg[*fabp10*:mcherry] background treated with DMSO or bafilomycin. Error bars indicate SEM. *P < 0.05. (E) Lipids staining of the liver in *larsb-I451F* larvae at 5 dpf treated with DMSO or bafilomycin. Scale bar: 20 μm. (F) Quantification of the lipid area in *larsb-I451F* larvae liver at 5 dpf treated with DMSO or bafilomycin. Error bars indicate SEM. **P < 0.01. Dpf: days post fertilization.

## Discussion

In this study, we demonstrated the pathogenesis of ALF in ILFS1 by excessive autophagy during Lar dysfunction. Liver dysfunction was most prominent in ILFS1 patients during infancy, which aligns with the finding of this study that *larsb-I451F* zebrafish exhibited liver abnormalities during the developmental stage. A histopathological analysis of *larsb-I451F* zebrafish showed the accumulation of lipid droplets in the liver, which mimicked the liver of ILFS1 patients caused by biallelic variants of the human *LARS* gene. In addition, enhanced autophagy was observed in the liver of *larsb-I451F* zebrafish. Inhibition of DGAT1, which converts fatty acids to triacylglycerols, improves lipid droplets in the liver of *larsb-I451F* zebrafish. Furthermore, the inactivation of autophagy by bafilomycin treatment significantly decreased the accumulation of intrahepatic lipids. These results suggest that Lars dysfunction in ILFS1 induced steatosis in the developing zebrafish liver via enhanced autophagy, pointing to the potential treatment of ALF by inhibiting autophagy.

In our previous study, *larsb*-knockout zebrafish exhibited progressive liver failure, anemia, and neurological defects that resembled the symptoms of human ILFS1 patients [29]. However, the liver of *larsb*-knockout zebrafish exhibited cytoplasmic loss due to severe damage, and early lethality precluded a further histological examination [29]. In the present study, we demonstrated the accumulation of lipids through enhanced autophagy in the liver of *larsb-I451F* zebrafish larvae. Although *larsb-I451F* zebrafish had the same amount of Lars protein as *larsb-WT* zebrafish, pathological variants of *LARS* led to a reduction in the aminoacylation activity of Lars, as previously reported in fibroblasts from ILFS1 patients [22]. The process of aminoacylation is executed through the precise functioning of leucine sensing and binding to the Lars protein, ATP binding, and structural alterations in Lars [45, 46]. Pathogenic variants of LARS1 that exhibit abnormalities in any of these functions lose their capacity to stimulate the mTORC1 pathway, which regulates autophagy [47]. Autophagy serves as an alternative energy source during nutrient deficiency by facilitating the breakdown of cellular components to produce fatty acids [48, 49]. However, excessive enhancement of autophagy beyond physiological limits can lead to autophagic cell death [50], which has also been confirmed in *larsb*-knockout zebrafish. While moderate autophagy serves as a protective mechanism against cell death during starvation, the surplus fatty acids generated during this process can be toxic and need to be directed into the mitochondria and used for energy production or stored as lipid droplets through DGAT1-mediated pathways [44, 51], as shown by *larsb-I451F* zebrafish in this study. The dysregulation of liver autophagy might differ between cases with complete deficiency and a partially retained function of Lars. Notably, the C-terminal region is known to interact with mTORC1 at the lysosomal membrane [26]. Further analyses using knock-in zebrafish with other genotypes will elucidate the mechanism by which Lar dysfunction activates autophagy.

*Larsb-I451F* zebrafish exhibited an atypical liver morphology at 5 dpf. In zebrafish embryogenesis, critical organ systems, such as the liver, rapidly develop by 5 dpf [52]. During this process, the complex mechanism of autophagy plays a crucial role in the regulation of cellular proliferation and differentiation. In zebrafish embryo development, autophagic activity sufficiently increases from 3 dpf to 5 dpf.[53, 54] In patients diagnosed with ILFS1, ALF is predominantly observed in the neonatal and infantile phases [22]. Given the resemblance between clinical liver pathology in ILFS1 patients and histopathological findings in *larsb-I451F* zebrafish larvae, it is plausible that liver damage is predominantly observed in neonates and infants due to defects in the *LARS* gene caused by increased autophagy. We further postulate that if remarkable and specific stimuli activate autophagy in cells, organ-specific damage can occur at any time during the lifespan.

Our findings suggest that dysregulation of autophagy caused by biallelic pathogenic variants of *larsb* leads to liver steatosis. Since significant similarities were observed between the liver tissues of human ILFS 1 and those of *larsb-I451F* zebrafish, this knock-in zebrafish more closely replicates ILFS1 than the *larsb*-knockout zebrafish. While patients with ILFS1 have a reduced risk of ALF after infancy, neurological and hematopoietic complications may relapse or newly appear in the long term. Unlike *larsb*-knockout larvae, *larsb-I451F* larvae can survive for a long time as adult zebrafish, so a straightforward evaluation of neurodevelopment and hematopoiesis can be achieved. Inborn errors of metabolism, such as Niemann-Pick disease type C and Gaucher disease, are known to present with distinct hepatic abnormalities during infancy and neurological symptoms in adolescence or adulthood. Similarly, citrin deficiency, which causes transient cholestatic liver disease in infancy, suddenly manifests as hyperammonemia in later adulthood after a long asymptomatic period. Consequently, long-term clinical trajectories can only be elucidated using model organisms capable of long-term observation. Previous case reports of ILFS1 are limited in number, and the long-term clinical characteristics of the surviving cases remain unclear. Zebrafish offer advantages as a suitable model organism for such observations and screening for potential drugs or chemical compounds.

*Larsb*-I451F zebrafish may thus serve as an optimal model for the long-term study of ILFS1 and may provide invaluable findings for further basic and clinical research.

## Materials and Methods

### Ethics statement

This study using human data was approved by the ethics committee of the Institutional Review Board of Oita University Hospital, Japan (approval no. 2565). Written informed consent was obtained from all participants. The animal study protocol was approved by the Institutional Review Board of Oita University (approval nos. 230501 and 4-5).

### WES analysis

Genomic DNA was extracted from the peripheral blood of the proband and his sister, and the parents were sequenced by WES. The sequence library was prepared using a Human All Exon V6 Kit (Agilent Technologies, Santa Clara, CA, USA) and sequenced using a 2500 Illumina with 125-bp paired-end reads (Illumina, San Diego, CA, USA). Sequence reads were aligned to GRCh38 and annotated using CompStor NOVOS and CompStor Insight (OmniTier, San Jose, CA, USA). First, variants with allele frequencies greater than 0.01 in gnomAD, 14 KJPN (jMORP), and our in-house exome variant data were removed. Next, the variants were narrowed down based on assumed modes of inheritance, such as autosomal dominant, autosomal recessive, X-linked, and compound heterozygous inheritance. Finally, three variants were segregated, one of which was inconsistent with the clinical symptoms (S1 Table) [30]. No pathogenic copy number variation was detected in the WES data. The two *LARS1* variants were confirmed by Sanger sequencing (ABI3130) using the primers 5′-GGGTCTCATAACAATGAATACTTC-3′ and 5′-GGGAAAAGGTAGGCTACAAGG -3’ for NM_020117:c.601T>G, and 5′-GGCAGTGTCGTAATGACATATAC-3′ and 5′-CCATAGAGATTCCTAGAGGG-3′ for c.1351A>T.

### Zebrafish maintenance

The zebrafish AB genetic background *larsb* mutant and Tg[*fabp10*:mcherry] were raised and maintained following standard procedures [34, 35]. They were maintained at 28– 29 °C under a 14-h:10-h light:dark cycle. Embryos were collected and housed at 28.5 °C.

All animal experimental procedures were performed in accordance with institutional and national guidelines and regulations. The study was conducted in compliance with the ARRIVE guidelines.

### Generation of the larsb I451F zebrafish line

The *larsb I451F* zebrafish line was generated via CRISPR/ Cas9 gene editing [55, 56]. The site of the *larsb* sgRNA target was 5′-CCAAAGCCAGAATGACAGAGAGA-3′ in the editing domain of the LARS protein. Single-stranded oligodeoxynucleotides (ssODNs) were designed with the following sequences (phosphorothioate modifications in the first and last nucleotides) and ordered as ultramers from Integrated DNA Technologies (Coralville, IA, USA) to generate single nucleotide polymorphism mutations: A*G*TGGCTTATTGGTTTGTTCTACCAGGTTCCCATCATTGAAATTCCAGGGTATGG GAATCTGTCAGCTCCACTGGTGTGCGATGAACTGAAGTTTCAAAGCCAGAATGAC AGAGAGAAACTGGCCGAGG*C*T. Cas9 protein (300 pg), gRNA (30 pg), and ssODNs (41 pg) were injected into one-cell-stage wild-type embryos. Mutations at the target site were verified using Sanger sequencing.

### Generation of transgenic zebrafish

Tg[*fabp10*:mCherry] fish expressing mCherry exclusively in hepatocytes were generated using the MultiSite Gateway kit (Thermo Fisher Scientific, Waltham, MA, USA) to produce vectors with Tol2 transposon sites [57]. A 2.8-kb promoter of the *fabp10* gene [34] was cloned into the p5E-mcs vector. Multisite Gateway cloning [58] was performed using the destination vector pDestTol2pA2, the 5′ entry vector containing the fabp10 promoter, the middle entry vector containing pME-mCherry, and the 3′ entry vector containing p3E-polyA. DNA constructs (25 pg) and Tol2 mRNA (25 pg) were injected into wild-type zebrafish embryos at the single-cell stage.

### Western blotting

Samples for Western blotting were lysed with lysis buffer (0.5% NP-40, 10% glycerin, 50 mM HEPES–KOH [pH 7.8], 150 mM NaCl, and 1 mM EDTA) using protease and a phosphatase inhibitor cocktail (Thermo Fisher Scientific). Protein samples were separated by capillary electrophoresis using 12-to 230-kDa Wes Separation Module capillary cartridges in the Simple Protein Wes system (ProteinSimple Wes; ProteinSimple. San Jose, CA, USA) according to the manufacturer’s protocol. The antibodies used were as follows: Lars (#13868; Cell Signaling Technology, Beverly, MA, USA; 1:50) and β-actin (A3854; Sigma-Aldrich, St. Louis, MO, USA; 1:100). The anti-rabbit and anti-mouse modules for the Wes kit (DM-001 and DM-002, ProteinSimple), which includes luminol-S, peroxide, antibody diluent 2, streptavidin-HRP, anti-rabbit secondary antibody, and anti-mouse secondary antibody, were used for detection. The intensities of the acquired chemiluminescence signals were quantified using the AlphaView and Compass software programs (ProteinSimple).

### Morphological analyses

Zebrafish larvae were placed in 3% methylcellulose, and images were acquired using a Leica M205 FA fluorescent stereo microscope (Leica, Wetzlar, Germany). Liver circularity was measured manually using the ImageJ Fiji software program (1.53t; National Institutes of Health, Bethesda, MD, USA).

### Histopathological staining and fluorescent immunostaining

Histopathological staining and fluorescent immunostaining were performed on the paraffin or frozen sections. For histopathological staining, samples were initially stained with hematoxylin solution for 20 s and rinsed with deionized water. They were then stained with eosin solution for 60 s and rinsed again with deionized water, and then they were dehydrated using a series of ascending ethanol concentrations. The excess blot was removed using xylene for 30 s (three repetitions). Finally, the coverslips were mounted using a mounting medium. The immunofluorescence analysis was performed using the following primary antibodies: anti-p62 (PM045; Medical & Biological Laboratories, Nagoya, Japan) and anti-LC-3 pAb (PM036; Medical & Biological Laboratories). Alexa Fluor 488 donkey anti-rabbit IgG (A21206; Molecular Probes, Eugene, OR, USA; 1:500) was used as the secondary antibody. Images were captured using a laser scanning microscope (BZ-9000; Keyence, Osaka, Japan).

### Fluorescent staining of accumulated lipids

Frozen samples were rinsed with phosphate-buffered saline. The samples were then stained with 1 μM Lipi Dye II solution (Funakoshi, Tokyo, Japan) in phosphate-buffered saline and incubated for 1 h at 37 °C. The cells were rinsed three times with phosphate-buffered saline and mounted with a fluorescence mounting medium (S3023; Dako, Agilent Technologies). Images were captured using a laser scanning microscope (BZ-9000; Keyence).

### Bafilomycin A1 and A922500 treatments

Embryos were treated with bafilomycin A1 (2.5 nM; EMD Millipore, Darmstadt, Germany), A922500 (2 mM; Sigma-Aldrich, St. Louis, MO, USA), or dimethyl sulfoxide (DMSO) as the control, in embryo medium from 72 to 120 hpf for morphological experiments. Water containing the drug was replaced daily.

### Statistical analyses

Statistical analyses were performed using the GraphPad Prism software version 8 (GraphPad Software, Inc., San Diego, CA, USA). All values are expressed as the mean ± standard error of the mean. Shapiro–Wilk and Brown–Forsythe tests were performed to analyze the normal distribution and homogeneity of the data, respectively. The different groups were compared using nonparametric independent samples Kruskal–Wallis test for non-normally distributed variables, and the results obtained were expressed as median and interquartile ranges. In contrast, when the data had a normal distribution, they were analyzed through a one-way analysis of variance (ANOVA) followed by Tukey’s pairwise comparison tests. Statistical differences in survival curves were analyzed using the log-rank (Mantel-Cox) test. Statistical significance was set at P < 0.05.

## Acknowledgements

We thank Sayaka Kai, Miki Nakamura-Ota, and Kaori Miura for their technical assistance.

## Funding Statement

Masanori Inoue was supported by the Japan Society for the Promotion of Science (22K15947) and the Kawano Masanori Memorial Public Interest Incorporated Foundation for the Promotion of Pediatrics. Toshikatsu Hanada was supported by the Japan Society for the Promotion of Science (20H03644), the Takeda Science Foundation, Kamizono Kids Clinic, and Mizoguchi Urology Clinic.

## Competing Interests

The authors have declared that no competing interests exist.

## Author Contributions

M.I. and M.M. collected the clinical data. K.Y. and T.K. performed the genetic analysis. M.I. generated mutant zebrafish and performed zebrafish phenotyping with the assistance of N.S. and H.S. M.I., W.S., S.S., and H.M. performed the histological analysis. R.H. provided key reagents and technical assistance for the generation of mutant zebrafish. M.I. drafted the manuscript. K.I. and T.H. coordinated the project, and reviewed and edited the manuscript. All authors read, revised, and approved the final draft.

## Supporting information

**S1 Table.**
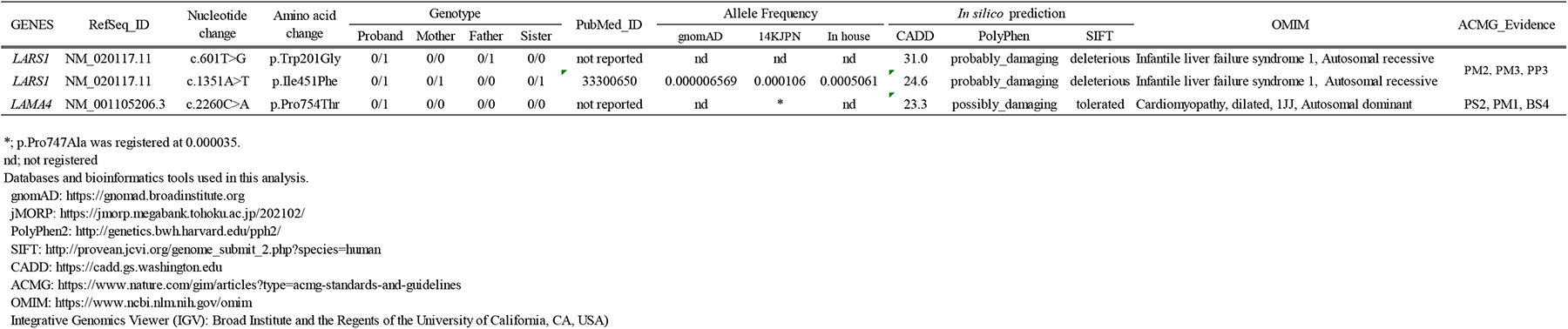
Segregated variants in the family.

**S1 Fig.**
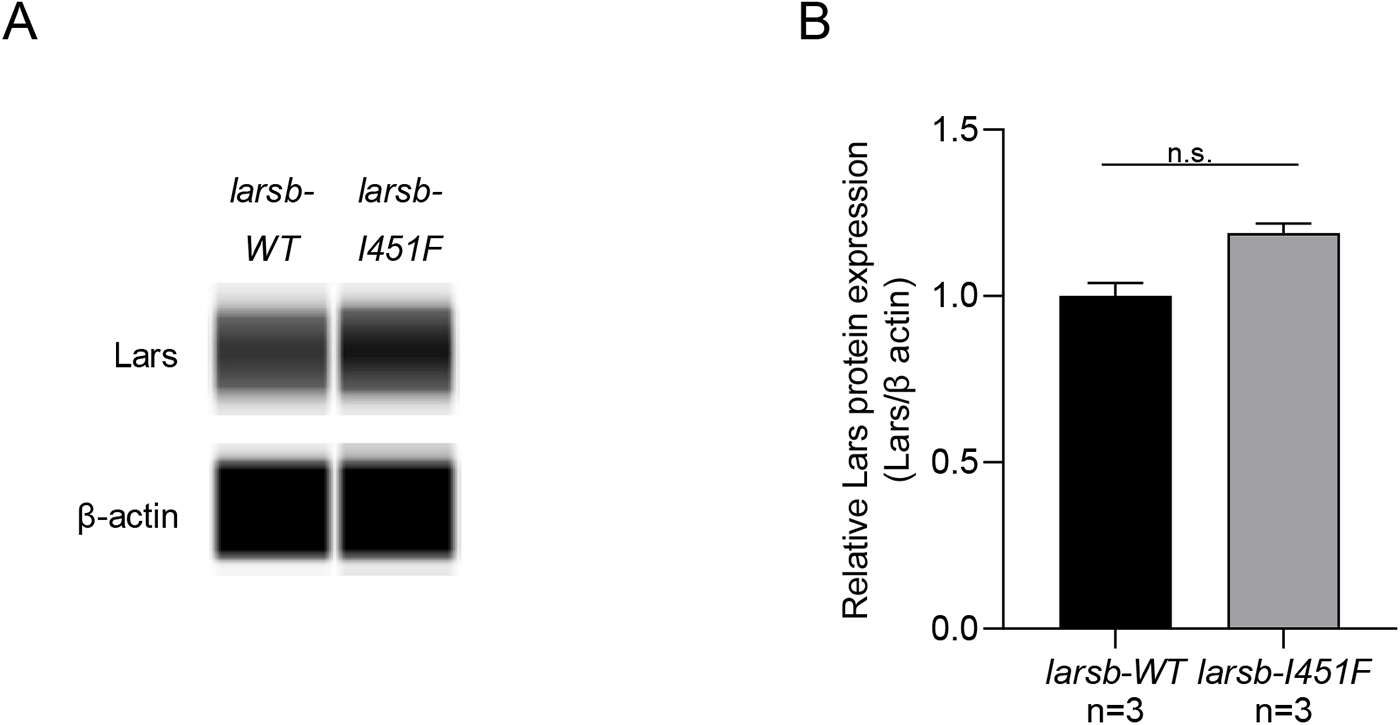
Western blot analysis of the Larsb protein expression in *larsb*-knockin zebrafish. (A) A Western blot analysis for Larsb protein in wild-type and *larsb*-knockin (*larsb-I451F*) zebrafish at 5 dpf. β-actin levels served as the loading control. (B) Densitometric quantification of the relative ratio of Larsb protein to β-actin protein in three independent experiments. Error bars indicate SEM. *P < 0.05. Dpf: days post fertilization.

**S2 Fig.**
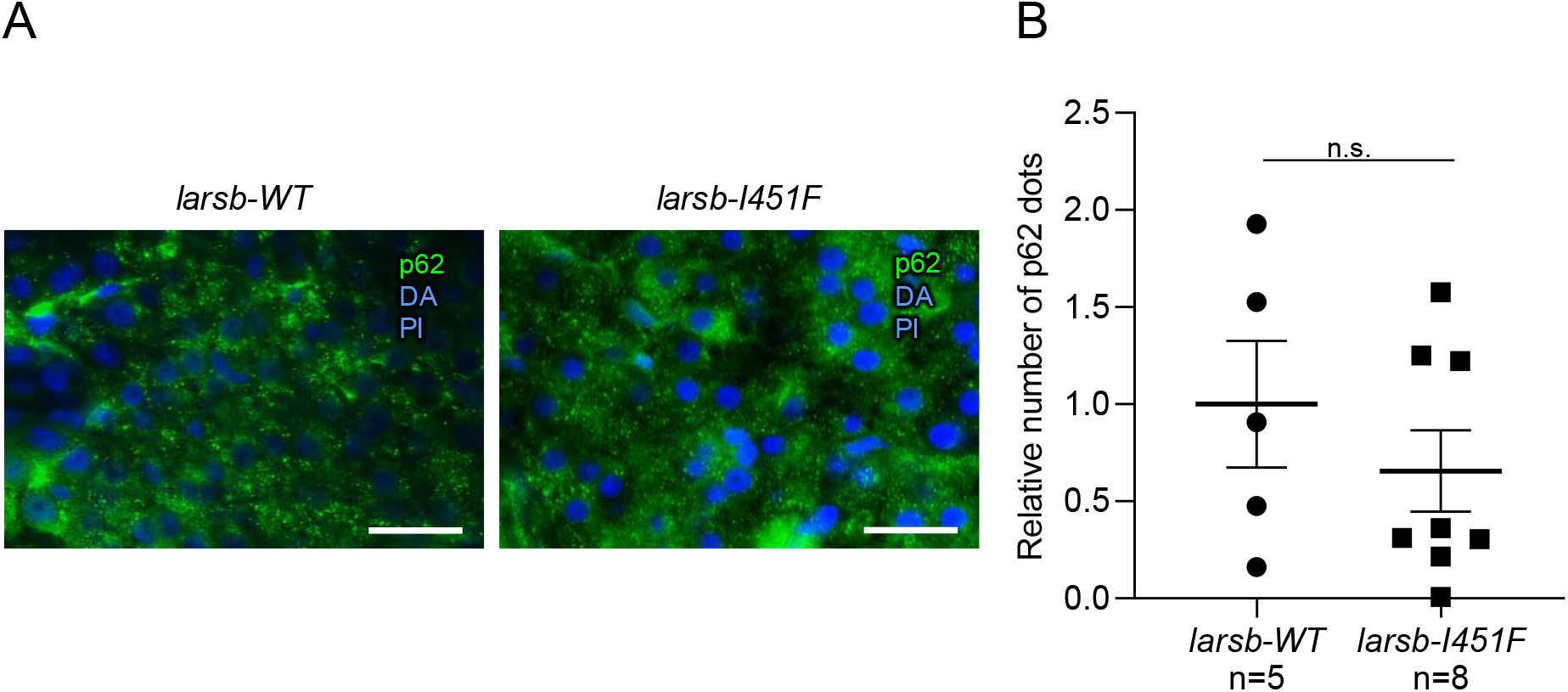
The evaluation of p62 in the liver of *larsb*-knockin larvae. (A) Immunostaining of p62 in the liver of *larsb*-knockin (*larsb-I451F*) larvae at 5 dpf. Scale bar: 20 μm. (B) Quantification of the number of p62 dots in *larsb-I451F* larvae liver at 5 dpf. Error bars indicate SEM. Dpf: days post fertilization.

**S3 Fig.**
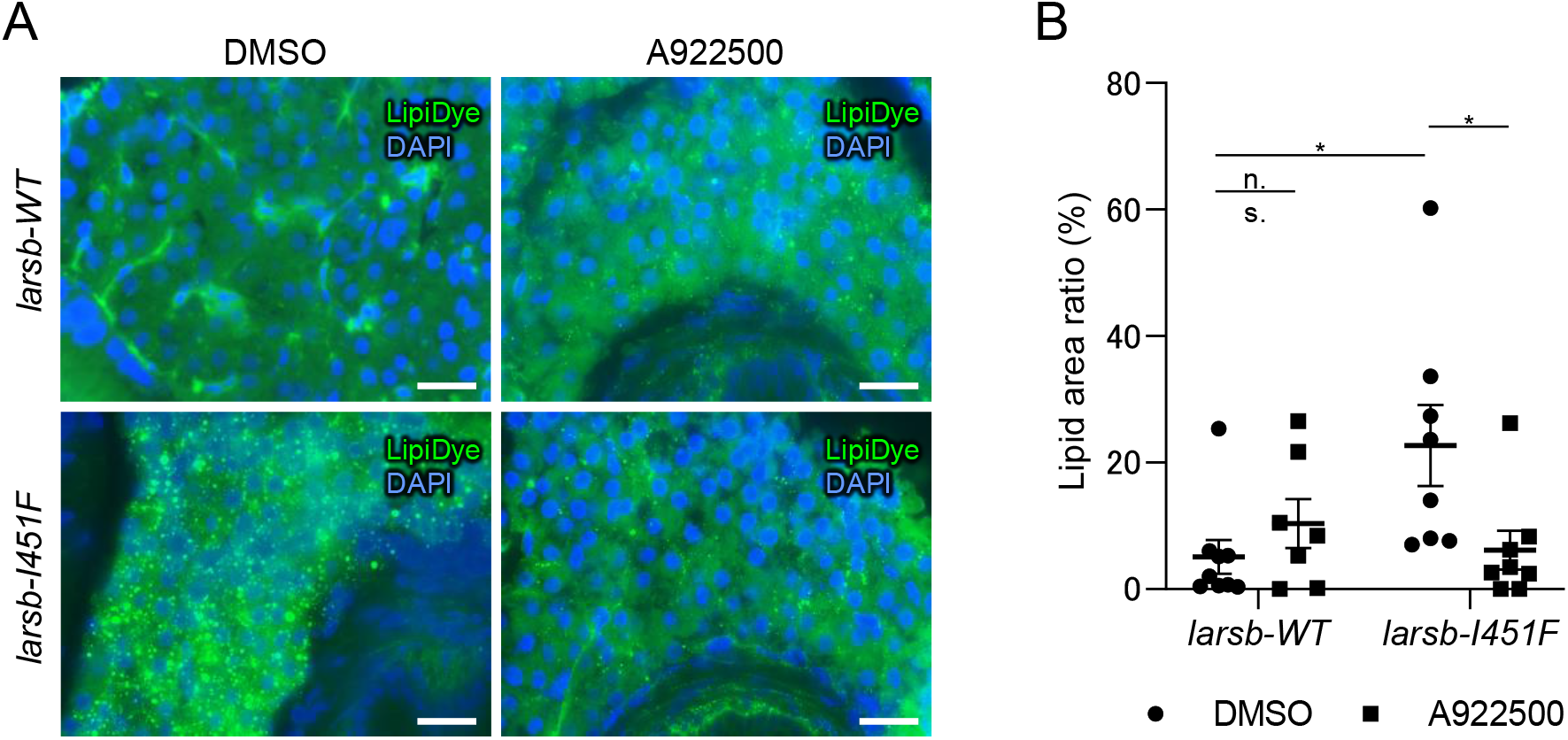
Inhibition of DGTA1 prevents the liver steatosis in *larsb*-knockin larvae. (B) Lipid staining of the liver in *larsb*-knockin (*larsb-I451F*) larvae at 5 dpf treated with DMSO or A922500. Scale bar: 20 μm. (B) Quantification of the lipid area in *larsb-I451F* larvae liver at 5 dpf treated with DMSO or A922500. Error bars indicate SEM. *P < 0.05. DGAT1: Diacylglycerol acyltransferase 1, dpf: days post fertilization.

## Notes

### Competing Interest Statement

The authors have declared no competing interest.

